# Microtubules sustain the fidelity of cellularization in a coenocytic relative of animals

**DOI:** 10.64898/2026.02.16.706138

**Authors:** Margarida Araújo, Marine Olivetta, Paolo Ronchi, Viola Oorschot, Arif Khan, Christian Tischer, Hiral Shah, Gautam Dey, Omaya Dudin

## Abstract

Cellularization is the coordinated division of a multinucleate cytoplasm into many cells.^1–3^ Multinucleation is a common life cycle strategy observed across eukaryotic lineages, including in microbial eukaryotes, fungi, plants and animals, and is associated with the ability to transition to a unicellular state through cellularization.^4^ In the best-studied model for this process, *Drosophila melanogaster*, cellularization requires the coordinated action of actin and microtubule (MT) networks to bring about the synchronous invagination of plasma membrane furrows, but the extent of conservation of these mechanisms across eukaryotes remains unknown.^1,5,6^ Here we investigate cellularization in the ichthyosporean *Sphaeroforma arctica*, a close relative of animals with a multinucleate life cycle stage.^7–9^ Using live cell imaging, ultrastructure expansion microscopy (U-ExM) and volume electron microscopy, we define the membrane, MT and actin dynamics that accompany cellularization in *S. arctica*. Using pharmacological inhibitors and centrifugation, we show that MTs, in addition to positioning nuclei, play a role in guiding nascent furrows to sustain equi-partitioning of nuclei and cytoplasm between daughter cells. Our findings indicate that cellularization is regulated through crosstalk between actin and MT networks, exhibiting mechanistic parallels with canonical cytokinesis, and establish *S. arctica* as a valuable model for investigating general principles of cellularization.

## Results & discussion

Multinucleate cells arise through two distinct mechanisms: syncytia that form by cell fusion, or coenocytes that result from nuclear division without cytokinesis.^2,4^ Some multinucleate cells persist throughout their lifetime, while others undergo cellularization, a specialized form of cytokinesis in which individual nuclei become encapsulated by the plasma membrane (PM) to form discrete cells.^1,2,4,10^ Cellularization occurs across diverse eukaryotic lineages, including insect embryos, the endosperm of flowering plants, chytrid fungi, and deep branching holozoans such as Ichthyosporea.^1,3,5,7,10–14^

In *Drosophila melanogaster*, cellularization during the 14th nuclear division transforms the syncytial blastoderm into an epithelial monolayer.^15,16^ This process depends on zygotic genome activation, nucleocytoplasmic ratio thresholds, and membrane expansion via Golgi-derived vesicles.^15,17,18^ Disruption leads to developmental defects, arrest or lethality.^1,19,20^ Actin and microtubules (MTs) contribute in system-specific ways: in *D. melanogaster*, actin drives furrow ingression, and MTs mediate nuclear positioning and vesicle transport.^5,6,21,22^ Similarly, endosperm cellularization in *Arabidopsis thaliana* relies on radial MTs and actin cables to maintain nuclear spacing and furrow position.^3,11^ In chytrid fungi such as *Spizellomyces punctatus*, cellularization occurs within a spherical syncytium containing dispersed nuclei.^10,23^ Centrosomes migrate to the cortex, initiating vesicle-associated membrane invaginations, while actomyosin networks drive three-dimensional furrow ingression to generate a polyhedral cellularization.^10^ Although MTs are dispensable for furrow formation, they are required for nuclear patterning and ciliogenesis.^10^ Across systems, uniform nuclear spacing and positioning are consistently associated with successful cellularization, underscoring a conserved geometric constraint governing this process.^3,10,15,24,25^

Among ichthyosporeans (Figure 1A), a lineage of close animal relatives, *Sphaeroforma arctica* has emerged as a tractable model for investigating mechanisms orchestrating cellularization.^7–9,26–28^ *S. arctica* exhibits a life cycle in which cytokinesis is uncoupled from mitosis: cells undergo multiple rounds of nuclear division without cell division, forming multinucleated coenocytes (Figure 1B).^7^ During mitosis, *S. arctica* undergoes closed mitosis driven by an acentriolar microtubule organizing center (MTOC) that bears similarities to the cell-cycle-dependent organization of fungal spindle pole bodies.^29^ Following coenocytic growth, *S. arctica* undergoes actomyosin-dependent cellularization through coordinated plasma membrane invaginations that progress inward from the cortex, forming a transient layer of polarized cells (Figure 1B and 1C).^7^ Although the invaginations appear to create distinct cellular compartments surrounding individual nuclei, if and how these compartments fully seal to complete cytokinesis remains unclear. Recent work has shown that this cellularization process is regulated by the nucleocytoplasmic (N/C) ratio, with cellularization timing tightly coupled to nuclear content relative to cytoplasmic volume.^27^ This developmental transition shares features with insect cellularization, including membrane furrow formation, use of conserved cytoskeletal regulators such as formins, Arp2/3, and myosin II, and transcriptional waves of cytoskeletal and adhesion genes.^6,7,21^

**Figure 1.**
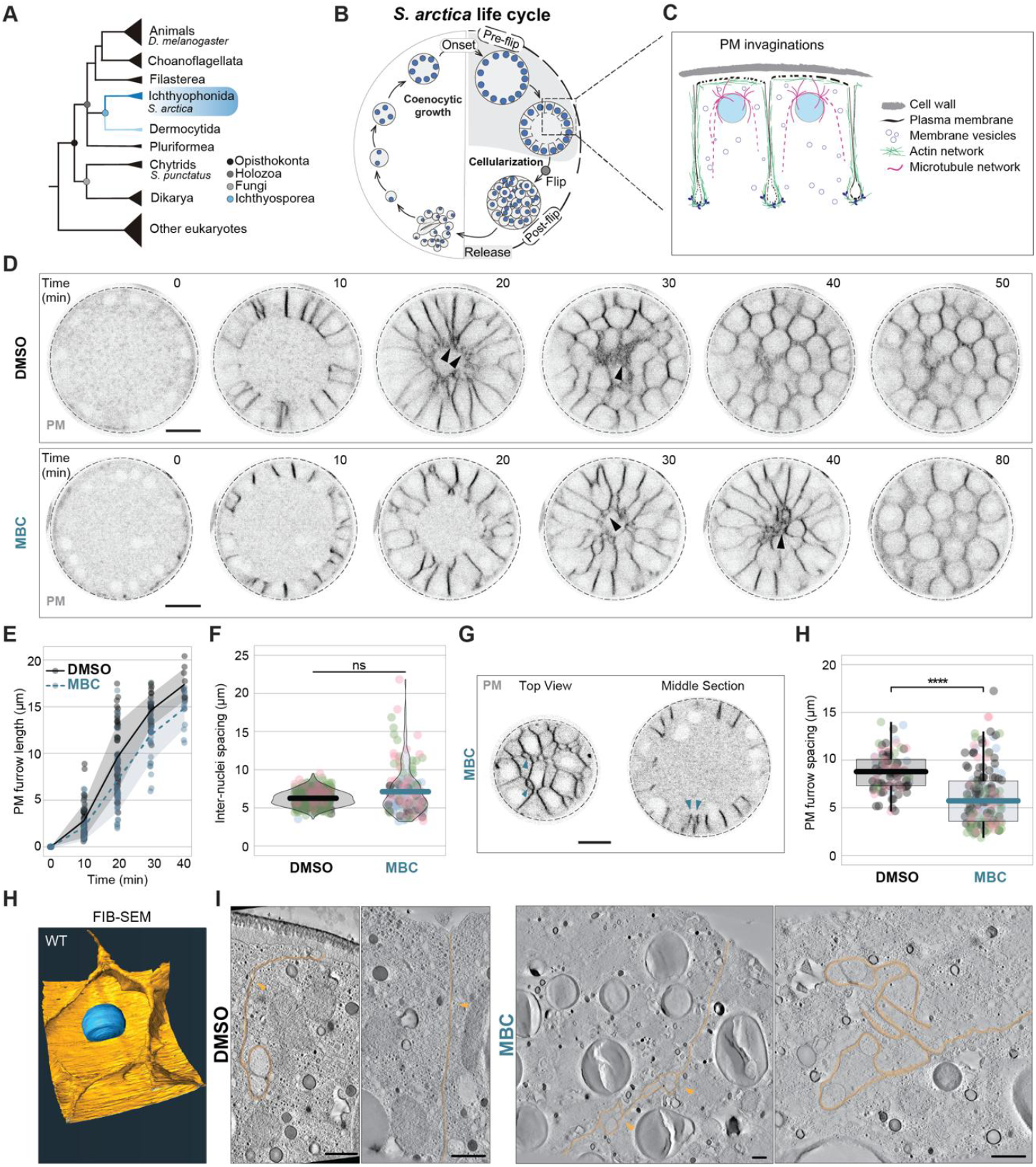
Loss of MTs alters cellularization dynamics. (A) Cladogram showing the position of Ichthyosporea within Holozoa and Opisthokonts. (B) Lifecycle of *S. arctica* highlighting cellularization with a grey background. (C) Schematic of cellularization at the cell cortex. (D) Time-course of cellularization membrane dynamics in control and carbendazim (MBC) treated coenocytes labelled with FM4-64 and imaged at 10-minute intervals. Arrows indicate membrane dynamics at furrow edges. Scale bar, 10 µm. (E) PM furrow lengths measured over time through invagination. Mean PM length ± SD at t=10, 20, 30 and 40 min are 2.89±1.83, 9.90±3.69, 14.70±1.56 and 17.37±1.48 µm for the DMSO control and 1.94±1.20, 6.53±3.33, 11.60±2.92 and 14.43±1.80 µm for the MBC treated cells, n = 14 coenocytes; analysis of variance (ANOVA) testing mean PM length for each treatment by time point: treatment p-value=0.042*, time point p-value=0.000234***. (F) Inter-nuclear spacing in control and MBC-treated cells. (G) Top and middle section of furrow spacing in MBC treated cells labelled with FM4-64. Scale bar, 10 µm. (H) Quantification of furrow spacing in control (n=31) and MBC-treated (n=15) coenocytes, p-value=4.5e−14****. (I) 3D segmentation of plasma membrane (yellow) and nucleus (blue) from a FIB-SEM volume of a cellularizing coenocyte. (J) TEM tomography slices showing furrow morphology (yellow outlines, arrowheads) in control and MBC-treated cells. Scale bar: 500nm.

In contrast, the contribution of MTs during the cellularization of *S. arctica* remains poorly understood. While MT depolymerization disrupts nuclear positioning and spacing, membrane invaginations still occur, suggesting that MTs are not required for furrow initiation.^7^ However, how MTs contribute to the spatial organization, timing, and coordination of cellularization, and how they interact with the actin cytoskeleton and membranes, remains unresolved.

To test the role of MTs in cellularization, we treated cellularizing coenocytes (34 hrs post-inoculation (hpi)) with the MT-depolymerizing agent carbendazim (MBC) for 2 hrs. As previously shown, unlike actin inhibition, MT depolymerization does not completely block cellularization in *S. arctica*, allowing us to monitor the altered dynamics in the presence of MBC.^7^ We used live-cell imaging with the lipophilic dye FM4-64 to monitor PM furrows and follow their ingression kinetics over time (Figure 1D). MT loss disrupted cellularization dynamics, delaying furrow closure by 10–15 minutes and producing uncoordinated ingression with variable rates, diagonal trajectories, and occasional bifurcations, in contrast to the uniform, perpendicular furrows observed in controls (Figure 1D). (Figure 1D, Video S1). Second, furrow ingression rates were reduced, progressing at 0.421 μm/min in MBC-treated cells versus 0.487 μm/min in controls with extensive variability (Figure 1E). Third, MBC-treated coenocytes failed to extend PM furrows to the same length as controls, achieving only 14.43 ± 1.80 μm compared to 17.27 ± 1.48 μm (Figure 1E). Notably, these invagination rates in *S. arctica* are comparable to those reported during *Drosophila* embryo cellularization, where slow and fast phase invagination proceed at 0.2 ± 0.02 μm/min and 1.2 ± 0.10 μm/min, respectively, to achieve a final furrow length of ∼35 μm.^30^

Beyond furrow kinetics, MT depolymerization severely disrupted the spatial organization of cellularization. While mean inter-nuclear spacing was not significantly different between conditions, MBC-treated coenocytes exhibited dramatically increased variability in nuclear positioning, with nuclei ranging from properly positioned to severely mislocalized (Figure 1F). Consistent with this, MBC treatment disrupted the regular spacing of PM furrows observed in controls. In control coenocytes, furrows were evenly distributed, enclosing one nucleus per cellular compartment (Figure 1H). By contrast, MBC-treated coenocytes displayed significantly decreased and irregular furrow spacing, resulting in some compartments lacking nuclei entirely (Figure 1G & H). This irregular cellularization was also reflected in the variable cell sizes produced after cellularization in MBC-treated coenocytes, in stark contrast to the uniform cell size in controls (Video S1).

To partly assess the structural basis of these defects, we obtained volumetric ultrastructural data of PM furrow organization. We performed focused ion beam scanning electron microscopy (FIB-SEM) and transmission electron microscopy tomography (TEM tomography). FIB-SEM of a cellularizing coenocyte enabled three-dimensional segmentation and volumetric reconstruction of both the plasma membrane and nucleus of a newly forming cellular compartment (Figure 1I, Video S2). TEM tomography at higher resolution revealed that MBC treatment resulted in aberrant membrane architecture, with furrows exhibiting bifurcated and convoluted morphology and abnormal fusion at the base of invagination, compared to the smooth, organized structure of control furrows (Figure 1J). Together, the delayed and disorganized furrow progression, along with mispositioned nuclei and irregular compartmentalization, indicate that MTs maintain the spatiotemporal precision of cellularization in *S. arctica*. This coordination between actomyosing-dependent furrows and MT networks likely reflects a conserved cytoskeletal role in organizing cellular domains within a shared cytoplasm, paralleling mechanisms observed in early-stage insect embryos.

### Microtubules track plasma membrane invaginations during cellularization

To determine how MTs contribute to furrow positioning and organization, we examined their spatial relationship with the plasma membrane during cellularization using ultrastructure expansion microscopy (U-ExM) on cellularizing *S. arctica* coenocytes (Figure 2A). U-ExM enables efficient immunolabeling in cell-walled microbial eukaryotes by permeabilizing their protective cell layer during the expansion process, thereby improving antibody accessibility.^29,31,32^ Leveraging this approach, we applied the recently developed U-ExM-compatible HAK-actin probe together with antitubulin antibodies to simultaneously visualize both cytoskeletal networks throughout cellularization (Figure 2B-D, Video S3).^33^ This represents a significant advance over standard immunofluorescence protocols, which fail to label or resolve these structures in *S. arctica*.

**Figure 2.**
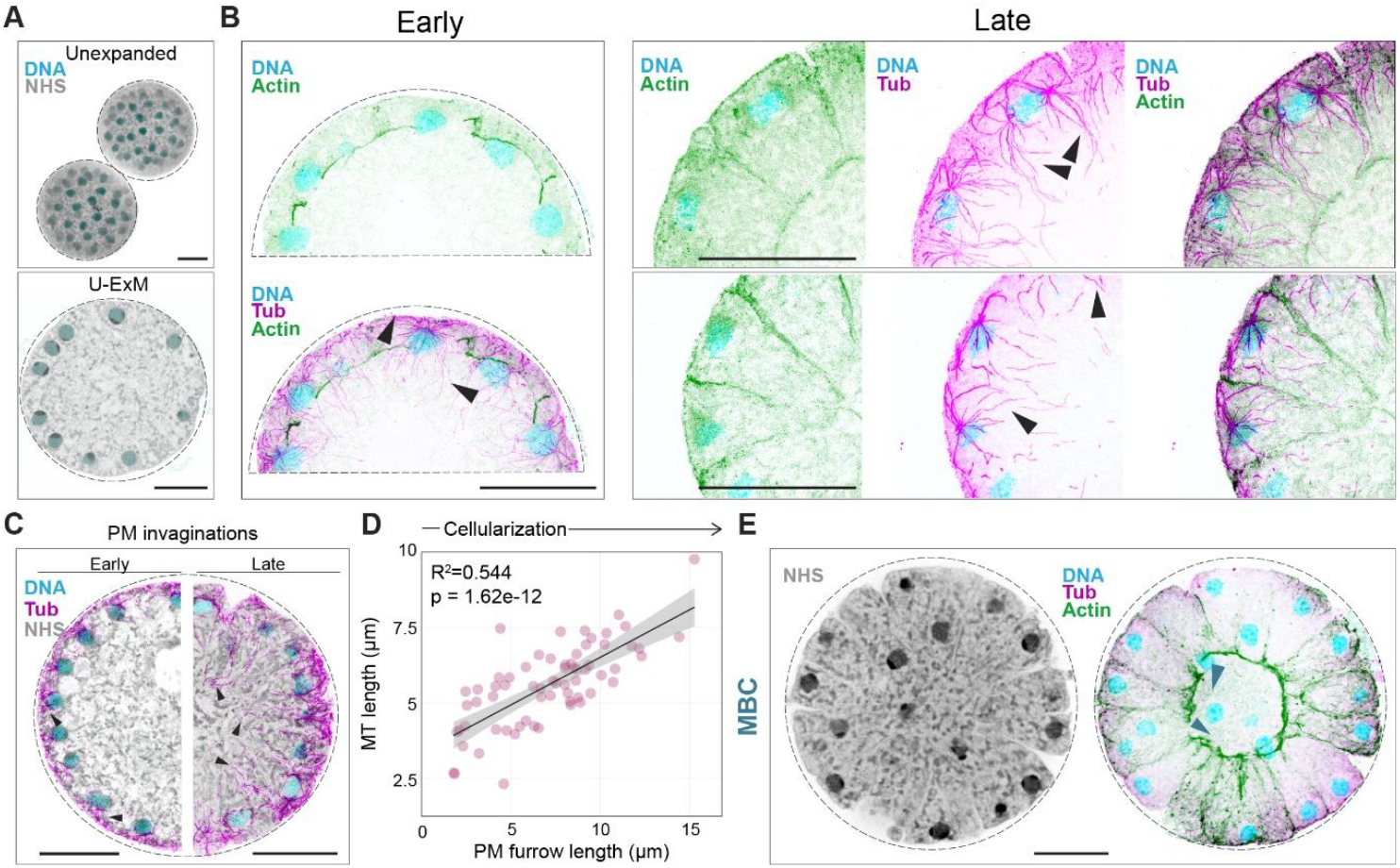
MT and actin cytoskeleton guide PM invagination. (A) Maximum intensity projections of non-expanded and expanded *S. arctica* coenocytes labelled for DNA (blue) and NHS-ester (grey). Maximum intensity projections of expanded cells labelled for actin (green), tubulin (magenta) and DNA (blue) at early (B) and late (C) stages of cellularization. Two examples of late cellularizing cells are showcased. Arrowheads mark cortical and furrow-tracking MTs. (D) MT organization at early and late stages of cellularization showcasing the extension of MTs length during cellularization. DNA (blue), NHS-ester (grey) and tubulin (magenta). Arrowheads indicate MTs. (E) MT lengths in relation to furrow length. (F) Maximum intensity projections of expanded cells labelled for actin, tubulin and DNA in MBC treated coenocytes. Scale bar 10 µm. Scale bars are adjusted for expansion factor.

At early stages, MTs extend from nucleus-associated (MTOCs) radially along nuclei and towards the cell cortex. As furrow formation progressed, we observed that MTs oriented towards the cell cortex and furrow-tracking MTs aligned along PM invaginations (Figure 2B & C, arrowheads). Actin localized at the base of membrane furrows throughout the invagination process. Although distinct contractile rings were not resolved, actin consistently accumulated at the invagination front. Notably, actin was also visible at the basal opening of nascent compartments that remained continuous with the shared cytoplasm (Figure 2C). In previous studies, actin accumulation has been interpreted as a marker of completed membrane sealing.^7^ However, our observations indicate that actin enrichment precedes final compartment closure, suggesting that membrane sealing occurs later than previously assumed. Consequently, the duration of the transient multicellular state in *S. arctica* might have been overestimated in earlier interpretations.

Quantitative analysis of segmented MT networks showed that cortical MT length scaled with furrow depth (R^2^ = 0.544, p = 1.62e−12), consistent with MTs elongating in coordination with plasma membrane invagination (Figure 2D & E). High-magnification views further supported that these bundles closely tracked the advancing furrow fronts, with longer MT extensions associated with deeper furrows during later stages (Figure 2D, arrowheads). While this correlation suggests a role for MTs in furrow progression, it remains unclear whether this involves active polymerization at the furrow front or utilization of pre-formed MT tracks. To further assess the requirement for MT dynamics in furrow organization, we analyzed MBC-treated coenocytes by U-ExM. Actin localization at the furrow base persisted, indicating that furrow formation can, as previously shown, proceed independently of MTs (Figure 2F). However, the spatial organization of furrows was severely affected, as shown in Figure 1. Furrows were often misaligned, and compartments frequently enclosed multiple nuclei or, conversely, lacked nuclei entirely. In several cases, nuclei were observed trailing between furrows or located beneath partially formed compartments, suggesting that improper nuclear positioning may interfere with furrow progression and sealing (Figure 2F).

These defects indicate that while furrow initiation is actin driven, MTs are required to position nuclei and align furrows accordingly. In their absence, cellularization proceeds but generates aberrant compartments with irregular geometry and compromised nuclear allocation. Taken together, these observations show that MTs in *S. arctica* reorganize during cellularization, transitioning from radial arrays around nuclei to furrow-aligned bundles that extend with membrane invagination. This dynamic reorganization, combined with actin accumulation at the furrow base, reveals a coordinated cytoskeletal architecture analogous to that observed in animal systems, suggesting that functionally analogous mechanisms regulate the spatial and temporal control of cellularization across distant eukaryotic lineages. While MBC treatment demonstrates that MT disruption leads to both nuclear misplacement and furrow disorganization, it remains unclear whether nuclei guide furrow positioning or whether forming furrows instead position the nuclei.

### Nuclear position guides furrow spacing

To assess the role of nuclear positioning in furrow organization, we centrifuged coenocytes undergoing cellularization to induce asymmetric displacement of nuclei along the cortex without disrupting MTs (Figure 3A).

**Figure 3.**
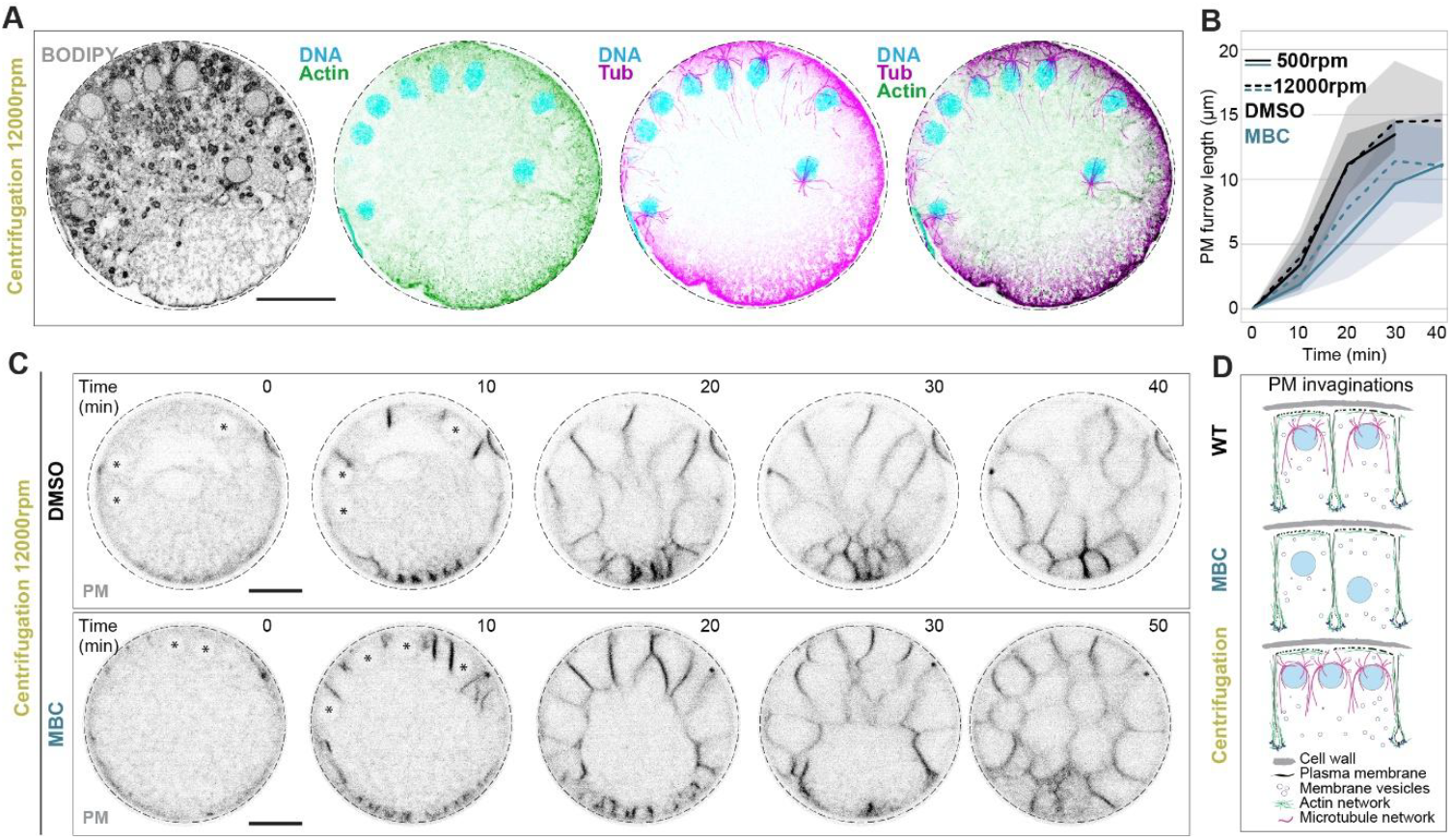
Nuclear positioning ensures fidelity of cellularization. (A) Maximum intensity projections of U-ExM images of coenocytes post centrifugation, labelled for membrane (grey), actin (green), tubulin (magenta) and DNA (blue). Scale bar 10 µm, adjusted for expansion factor. (B) PM furrow length over time in DMSO (black) and MBC (blue)-treated coenocytes post centrifugation. (C) Time lapse images showing PM furrow ingression at 10-minute intervals in centrifuged coenocytes treated with DMSO or MBC. Scale bar 10 µm, asterisks indicate nuclei. (D) Cartoon showing impact of perturbations (MBC and centrifugation) on nuclear and cytoskeletal organization during PM invagination.

This approach builds on previous work in which centrifugation was used to perturb the spatial relationship between nuclei and the cortex and demonstrate that cellularization in *S. arctica* is sensitive to nuclear position and local nucleocytoplasmic ratios.^27^ In our experiments, centrifugation resulted in nuclei accumulating on one side of the coenocyte, creating a strong spatial asymmetry in nuclear distribution (Figure 3A). U-ExM imaging, including BODIPY-ceramide labelling for enhanced membrane visualization, confirmed this asymmetric nuclear redistribution (Figure 3A).

Following centrifugation, furrow ingression proceeded with kinetics comparable to controls, confirming that the core machinery for membrane trafficking and actin-driven invagination remained functional (Figure 3B). However, furrow spatial patterning was dramatically altered: in nuclear-depleted cortical regions, furrow initiations were more frequent and closely clustered, but failed to progress deeply. In nuclear-enriched regions, furrows progressed with kinetics comparable to controls but were misaligned, frequently enclosing multiple nuclei per compartment rather than the single nucleus observed in controls (Figure 3C, Video S4). Despite the preservation of cortical MTs, the regular spacing of the cytoskeletal network was disrupted, and furrows no longer exhibited the uniform spacing characteristic of control coenocytes. These findings reveal that nuclear position directs the sites of furrow initiation and proper compartmentalization. The mechanism underlying this spatial control remains to be fully resolved: it may involve MT-mediated physical coupling between nuclei and cortical membrane domains, localized signals emanating from nucleus-associated MTOCs, local nucleocytoplasmic ratio effects, or a combination of these factors. The observation that furrows track nuclear position even when nuclei are artificially displaced indicates that nuclei (and their associated MTOCs) serve as spatial landmarks that pattern membrane invagination. Together with the MBC experiments demonstrating that MT depolymerization disrupts both nuclear spacing and furrow organization while furrow ingression itself persists, these results support a model in which the nucleus-MTOC unit functions as an organizational hub: MTs maintain nuclear distribution to establish properly spaced cortical domains where actin-driven membrane invagination proceeds with correct geometry and nuclear allocation (Figure 3D). However, while these experiments define how cellularization is spatially patterned, the cellular machinery driving membrane invagination itself remained to be identified.

### *S. arctica* cellularization relies on membrane trafficking

The previous experiments show that furrow invagination proceeds even when furrows are forcibly mispositioned. This dissociation indicates that the processes of new membrane addition and furrow positioning are controlled independently. To identify the cellular processes driving furrow invagination, we examined the role of membrane trafficking. In early *Drosophila* embryos, the membrane expands from a reservoir of microvilli localized at the apical cortex.^34,35^ The first phase of this membrane invagination is supplemented partly by Golgi-derived vesicles. These vesicles are transported in a MT-dependent manner and are stalled with colcemid or colchicine treatment which eventually prevented furrow invagination.^20,25,36^ It is unclear if such a membrane reservoir is present at the *S. arctica* cortex where the furrows are first initiated. Brefeldin A treatment in *Drosophila* embryos inhibits furrow progression in the final stage of cellularization.^17,20^ To test whether membrane trafficking contributes to furrow dynamics in *S. arctica*, we treated coenocytes with Brefeldin A (BfA), which inhibits protein transport from the endoplasmic reticulum (ER) to the Golgi apparatus. High-dose BfA treatment interferes with and in some cells even blocks cellularization, confirming that Golgi-mediated vesicle trafficking is required for cellularization (Supp. Figure 1). To examine the specific effects on furrow organization, we used live imaging to follow the dynamics of cellularization in *S. arctica* coenocytes upon mild membrane trafficking impairment in presence of BfA (Figure 4A).

**Figure 4.**
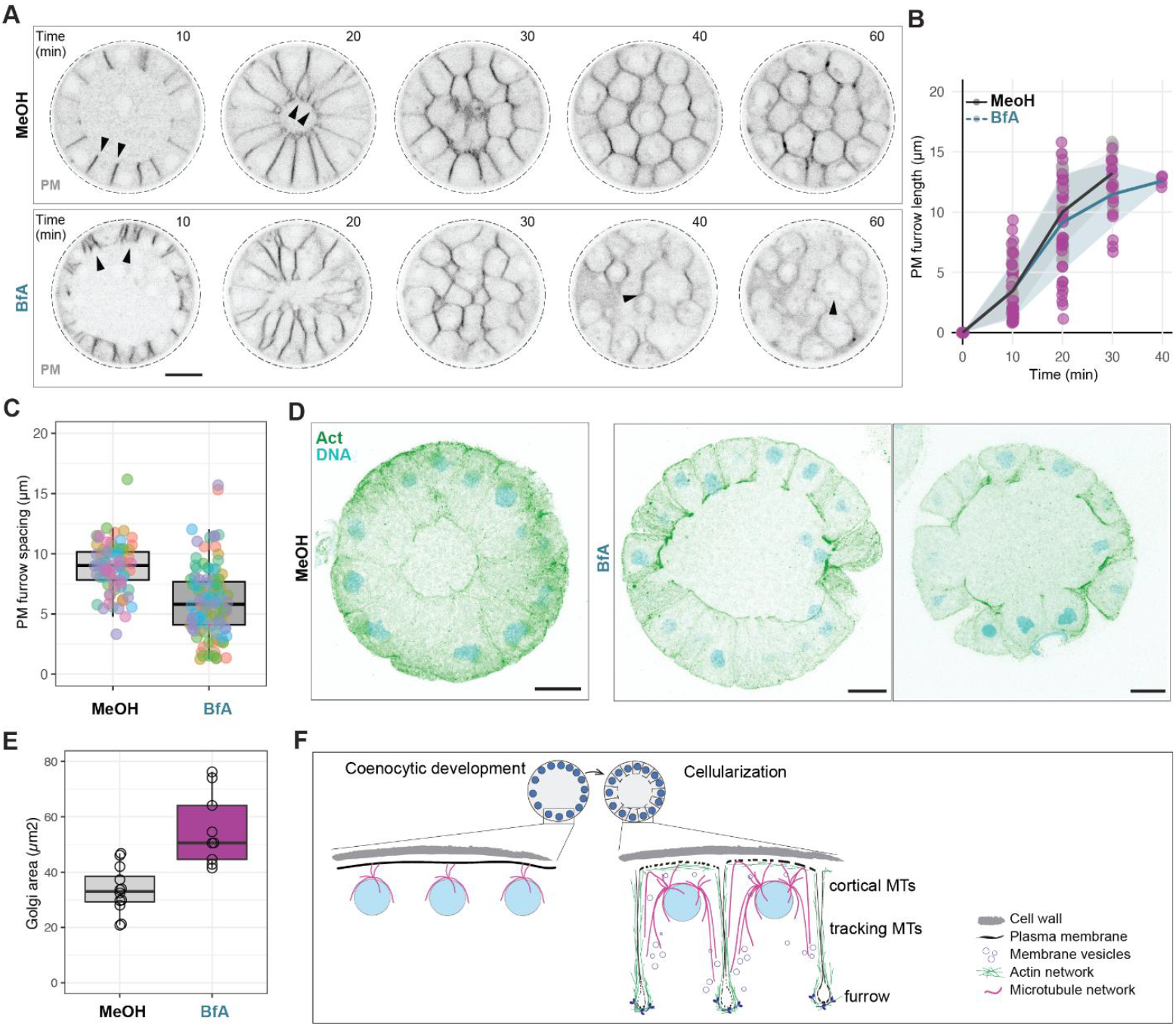
Brefeldin A inhibition leads to furrow mispositioning. (A) Membrane invasion dynamics in control (methanol) and Brefeldin A (BfA) treated coenocytes labeled with the FM4-64 membrane dye. Scale bar, 10 µm. (B) PM furrow length progression over time in methanol and BfA-treated coenocytes. (C) Quantification of PM furrow spacing for methanol-treated and BfA-treated coenocytes, n = 7 coenocytes. T-test p-value<0.0001****. (D) MIP of U-ExM images of control and BfA-treated cells labelled for DNA and actin. Scale bar 5 µm, adjusted for expansion factor. (E) Golgi area measured in control (methanol) and BfA-treated cells labelled with BODIPY ceramide. (F) Model showing MT and actin cytoskeleton guiding furrow ingression in cellularization.

Using this milder BfA treatment, we found that membrane furrows are still formed and can ingress at WT rates, however, they appear mispositioned like those seen upon MT depolymerization (Figure 4A & 4B). In agreement with time-lapse imaging, U-ExM images show misplaced furrows during cellularization when treated with BfA treatment (Figure 4C & 4D). Inhibition of membrane trafficking by 1.5 ug/ml BfA affected Golgi organization (Figure 4E and Supp. Figure 1B & 1C).

In summary, our results indicate that actin and MTs coordinate to guide membrane dynamics and spatial organization during cellularization in the ichthyosporean *S. arctica* (Figure 4F). Actin delineates furrow boundaries and accumulates at the invaginating front, while MTs dynamically reorganize to extend along furrows and maintain the regular spacing of nuclei and compartments. Disruption of either membrane trafficking or MT organization results in furrow misalignment and aberrant nuclear partitioning, uncoupling furrow initiation from spatial fidelity. Critically, cellularization operates through two coordinated modules: the nucleus-MTOC unit and its associated MTs establish spatial patterning, while actin and membrane trafficking drive the mechanics of furrow ingression. These pathways are separable yet must be coordinated for successful cellularization. The conservation of such cytoskeletal interplay in *S. arctica*, a close relative of animals, underscores the evolutionary depth of cellularization mechanisms and suggests that this modular architecture may represent a general principle enabling cellularization across diverse eukaryotic lineages and life cycle strategies.

## Acknowledgments

We thank the Gönczy, Dey and Dudin labs for discussions. We thank the Advanced Light Microscopy Facility and the Electron Microscopy Core Facility at the European Molecular Biology Laboratory (EMBL) for their support. H.S. was supported by the EMBL Interdisciplinary Postdoctoral Fellowship (EIPOD4) programme under Marie Sklodowska-Curie Actions Cofund (grant agreement no. 847543). G.D., M.A. and H.S. are supported by the European Union (ERC, KaryodynEvo, 101078291). G.D. and O.D. acknowledge the support of the EMBO Young Investigator Programme. M.A., P.R., V.O., A.K., C.T. and H.S. acknowledge core support from the European Molecular Biology Laboratory. M.O. and O.D., were supported by a Swiss National Science Foundation Starting Grant (TMSGI3_218007).

## Author contributions

Conceptualisation: M.A., O.D., G.D.; Methodology: M.A., M.O., P.R., V.O., A.K., C.T., G.D., O.D.; Validation: M.A., M.O., P.R., V.O., A.K., C.T., G.D., O.D.; Formal analysis: M.A., M.O., P.R., V.O., A.K., C.T., G.D., O.D.; Investigation: M.A., M.O., P.R., V.O., A.K., C.T., G.D., O.D.; Writing - Original Draft: M.A., H.S., G.D., O.D.; Writing – Review & Editing: M.A., H.S., M.O., P.R., V.O., A.K., C.T., G.D., O.D.; Visualisation: M.A., H.S., G.D., O.D.; Supervision: G.D., O.D.; Project administration: G.D., O.D.; Funding acquisition: G.D., O.D.

## Data availability

All image data related to this study are deposited at BioImage Archive 10.6019/S-BIAD2749. All quantification presented in the figures is shared as Data S1.

## Supplemental Information

**Supplementary figure 1:**
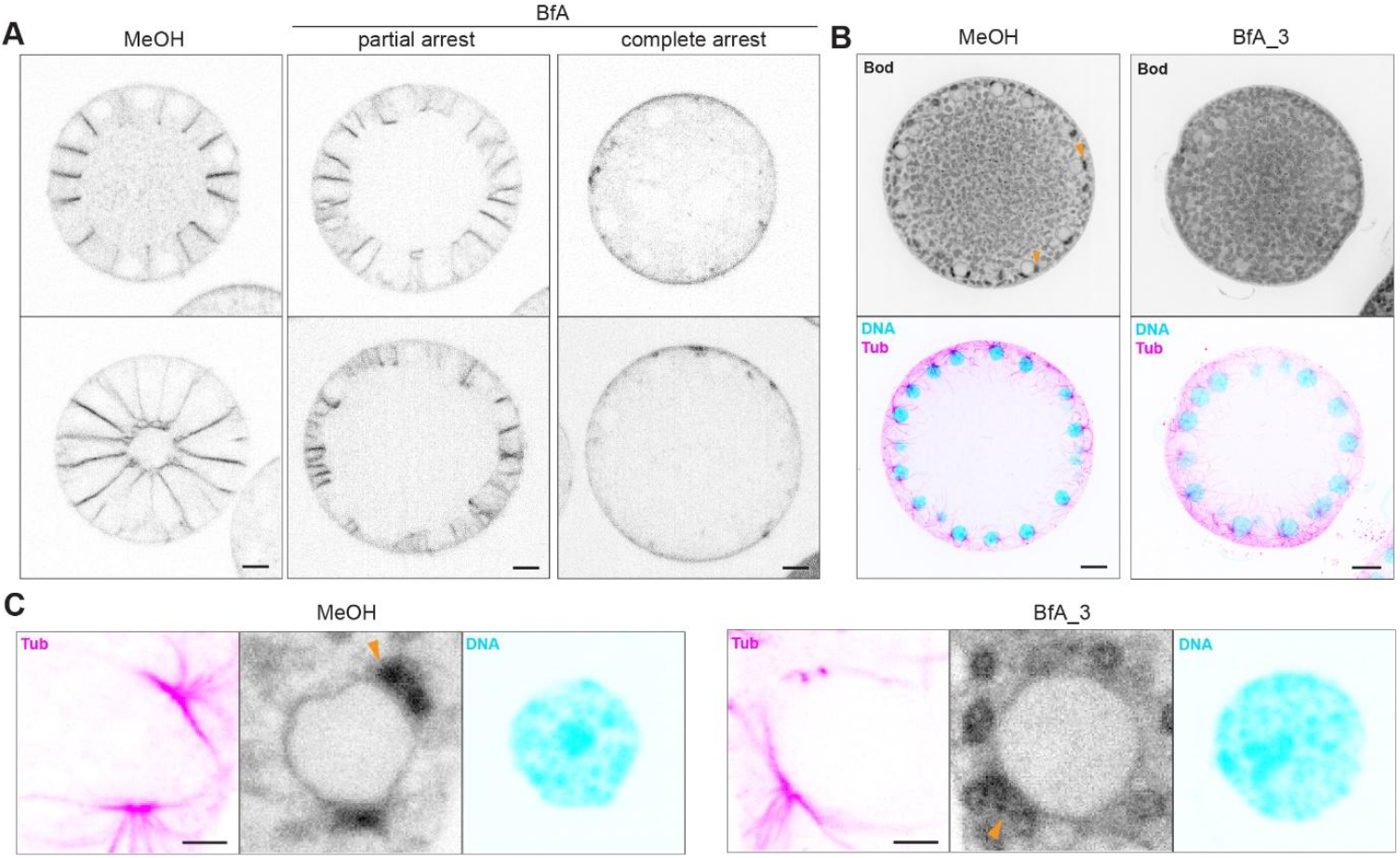
BfA treatment interferes with cellularization. (A) PM organization in methanol, 1.5 µg/ml BfA and 3µg/ml BfA-treated coenocytes labelled with FM4-64. The top and bottom panel shows two different coenocytes in each case. (B) Maximum intensity projections of U-ExM images of MeOH and 3µg/ml BfA-treated cells labelled for membrane, DNA and tubulin. Yellow arrowheads indicate nuclear-associated Golgi. Scale bar 5µm. (C) Zoom-in view of nuclei from cells shown in (B). Scale bar 1µm. Scale bars in U-ExM images are adjusted for expansion factor.

**Video S1:** Time-lapse movie showing cellularization in DMSO (left) and MBC-treated (right) coenocytes labelled with FM4-64 and imaged at 10-minute intervals. Scale bar 10 µm.

**Video S2:** FIB-SEM volume of cellularizing cell.

**Video S3:** U-ExM volume of cellularizing compartments. Tubulin (magenta), DNA (blue) and NHS-ester pan-labelling (grey).

**Video S4:** Time-lapse movie showing cellularization in control (500 rpm, left) and post centrifugation (12000 rpm, right) in MBC-treated coenocytes labelled with FM4-64 and imaged at 10-minute intervals. Scale bar 10 µm.

## Materials and Methods

### *S. arctica* culture and growth conditions

*Sphaeroforma arctica* was provided by the laboratory of Omaya Dudin. *S. arctica* cultures were maintained at 17°C in marine broth (Difco, 37.4 g l^−1^) in culture flasks (Falcon® 25cm^2^ Rectangular Canted Neck Cell Culture Flask with Vented Cap, Corning, product number 353109) and synchronized as previously described.^7,26,27^ Briefly, for synchronization, 1/16 marine broth (MB) was prepared by diluting MB in artificial seawater (Instant Ocean, 37 g l^−1^). Cultures were diluted 1:50 in 1/16 MB and grown for 5 days. 5 day old cultures were inoculated 1:60 - 1:70 in cold marine broth to get a synchronized culture. To obtain coenocytes undergoing cellularization, coenocytes were fixed within 35.5-38hrs hpi. For experiments, *S. arctica* cultures were diluted 1:200 every week, while for maintenance coenocytes were diluted 1:1000 every 2 weeks and restarted from a cryopreserved stock in 10% DMSO every 6 months.

### Effect of inhibitor treatments on *S. arctica* cellularization process

For microtubule perturbation during cellularization, coenocytes grown for 34hrs were treated with 25 μg ml^−1^ of carbendazim (MBC, Sigma 378674, using a 1:100 dilution from a freshly prepared stock solution at 2.5 mg ml^−1^ in DMSO for 2hrs). Cells were collected and fixed at 35.5, 36, 37 and 38 hpi. To inhibit membrane trafficking, coenocytes were treated with 1.5 μg ml^−1^ Brefeldin A (eBioscience™ Brefeldin A Solution 1000x in methanol, catalog number 00-4506) at 34 hpi.

### Chemical fixation and staging for cellularization by Phalloidin and Hoechst staining

Cell culture flasks were scraped and the suspension was collected into 15 ml Falcon tubes and centrifuged for 5 minutes at 500 rpm at 17°C. The supernatant was removed and coenocytes were transferred to 1.5 ml microfuge tubes and fixative was added for 30 min. The cells were fixed with 4% formaldehyde, 0.025 % Glutaraldehyde (Grade I, 25% in water, Sigma, G5882-50ML) in 250 mM sorbitol solution. The fixed cells were washed twice with 1× phosphate buffer saline (PBS) and resuspended in 100 μl of 1XPBS. For staining of actin and DNA, a staining solution containing Hoechst 33342 (20 mM, 1000x stock) at 1:500 and Alexa Fluor™ 568 or 488 Phalloidin (400x stock, A12380 and A12379, respectively, Invitrogen) at 1:200 dilution was prepared in 1x PBS. Briefly, 2 μl of this staining solution was placed on a glass slide and 2 μl of the fixed cell suspension was added and mixed into the staining solution, immediately covered with an 18×18 mm diameter coverslip and sealed with nail polish. Glass slides were incubated for 5 min at room temperature covered from light and imaged immediately after.

### Ultrastructure Expansion Microscopy

U-ExM was performed as previously described.^29,37^ Briefly, the cells were fixed as above. The fixed cells were then allowed to attach to 6 or 12 mm poly-D-lysine-coated (P1024-10MG, Sigma) coverslips for 1 h. For crosslinking with the HAK-Actin probe (JASP-HA, Spirochome), attached coenocytes were washed once with a drop of 1x PBS and then permeabilization with 1% PBS-T (Tween 20) for 30 min by adding a drop onto each coverslip.^33^ After permeabilization, coverslips were incubated with the HAK-Actin probe at 1:500 dilution in 1x PBS (15-20 μl per 6-mm coverslip) for 45 min at room temperature. Then, the HAK-Actin incubation solution was removed and coverslips were washed with 1x PBS. To crosslink the HAK-Actin probe, coenocytes were fixed again (4% Formaldehyde + 0.025% Glutaraldehyde + 250 mM sorbitol in PBS, freshly prepared) for 10 min at room temperature. The fixative solution was washed twice with 1x PBS. This was followed by anchoring in 1% acrylamide/0.7% formaldehyde solution overnight at 37 °C. A monomer solution (19% (wt/wt) sodium acrylate (Chem Cruz, AKSci 7446-81-3), 10% (wt/wt) acrylamide (Sigma-Aldrich A4058), 0.1% (wt/wt) N,N′-methylenebisacrylamide (Sigma-Aldrich M1533) in PBS) was used for gelation and gels were allowed to polymerize for 1 h at 37 °C in a moist chamber. For denaturation, gels were transferred to the denaturation buffer (50 mM Tris pH 9.0, 200 mM NaCl, 200 mM SDS, pH to 9.0) for 15 min at room temperature and then shifted to 95°C for 1 h. Following denaturation, expansion was performed with several washes in water. After expansion, gel diameter was measured and used to determine the expansion factor. For all U-ExM images, scale bars indicate actual size; rescaled for gel expansion factor.

For immunostaining, all antibodies were prepared in 3% BSA in 1x PBS with 0.1% Tween 20. Primary antibodies (alpha- and beta-tubulin antibodies, AA344 and AA345; ABCD antibodies) were used at 1:500, the anti-HA antibody (HA Tag Monoclonal Antibody (2-2.2.14, Catalog # 26183, Invitrogen) was used at 1:250, and incubated overnight at 37°C. This was followed by three washes for 10 min at room temperature and addition of the secondary antibody. Goat anti-mouse secondary antibody - Alexa Fluor 488 (Thermo A-11001) or 647 (A-21235) were used at 1:250, goat anti-guinea pig secondary antibody conjugated to Alexa Fluor 488 (Thermo A-11073) or 647 (Thermo A-21450), were used as secondary antibodies at 1:500. Pan labelling of U-ExM was done at 1:500 with Dylight 405 (ThermoFischer, 46400) or Alexa Fluor NHS-Ester 594 (ThermoFischer, A20004) in 1× PBS or NaHCO3 for 1.5-2 h or overnight (1:700 dilution). For membrane labelling, gels were stained with BODIPY TR ceramide (ThermoFischer D7540, 2 mM stock in DMSO) at 1:500 - 1:1000 dilution in 1× PBS for 2h at room temperature. DNA was stained with Hoechst 33352 at a final concentration of 0.4 μM.

### Light microscopy

Immunolabelled U-ExM gels were placed in poly-l-lysine-coated Ibidi chamber slides (two-well, Ibidi 80286; four-well, Ibidi 80426). Gels were cut to fit the Ibidi chambers and added onto the wells. The gels were overlaid with water to prevent drying or shrinkage during imaging. Gels were imaged using a Nikon-CSU-W1 Sora microscope with an Apo LWD 40x WI Lambda-S / 1.15 / 0,60 / water immersion objective with the lasers 405 nm (120 mW), 488 nm (200 mW), 561 nm (150 mW) and 638 nm (200 mW), or a Nikon A1 confocal microscope with a CFI Apo LWD Lambda-S 40x/1.15 WI 0,60 WD = 610 µm objective and LU-N4 405/488/561/640 laser unit. Z stacks were acquired with 130 (half coenocyte)-300 (whole coenocyte) slices using a Z step size of 0.5 μm and a zoom factor of 2.

### Live cell imaging

For live cell imaging, cells were synchronized as described above and grown for 34 hpi in marine broth at 17°C in 4-well ibidi chamber slides (Ibidi; 80426). To induce MT depolymerization and monitor cellularization dynamics, coenocytes were treated for 2h with either DMSO as a control or the MT inhibitor Carbendazim (MBC, Sigma Aldrich, 2.5 mg/ml stock in DMSO, final concentration 25 μl/ml). To perturb membrane trafficking during cellularization, coenocytes were treated for 2h with either methanol at 1:2000 dilution as a control or the membrane trafficking inhibitor BrefeldinA at 1.5 μg/ml or 3 μg/ml (1000x stock in methanol). The membrane dye FM4-64 (ThermoFisher; ref: T13320) was directly added to the medium 30 min before imaging at a final concentration of 5 μM (1:2000 dilution from a 5 mg/ml stock in DMSO). A Nikon-CSU-W1 Sora microscope with controlled temperature at 18°C was used for live cell imaging. Coenocytes were imaged with a 60x water immersion objective (SR P-Apochromat IR AC 60x WI/ 1.27) and a TRITC emission laser was used at 5% laser power with 50 ms exposure time. Z step size = 0.8 μm, 40 slices, 10 min time intervals.

### TEM tomography and tilt series reconstruction

For sample preparation for EM, coenocytes were diluted 1:50 in 1/16 marine broth and grown for 5 days to obtain saturated cultures. The saturated cultures were then diluted 1:70 in fresh cold marine broth (Falcon® 75 cm^2^ Rectangular Canted Neck Cell Culture Flask with Vented Cap, Corning, product number 353136) and grown for 28h. A volume of 20 ml per time point per condition was used. To increase synchronization, coenocytes were scraped from flasks, collected by centrifugation at 500 rpm at 17°C and washed 3 times in sea water to remove all the nutrients from the medium. Culture flasks were also washed with sea water, and cell suspensions in sea water were distributed with same initial volume into the culture flasks. Coenocytes were maintained in sea water for 8h and to enrich for cellularization events and treatment with DMSO (1:100) and MBC (2.5 mg/ml stock in DMSO, final concentration 25 μl/ml) was performed in the last 2 hours before high pressure freezing (HPF). HPF was done at time points 36 and 36.5h hpi. *S. arctica* cells were concentrated by centrifugation at 500 rpm for 5 min at 17°C and high-pressure frozen with the HPM010 (Abra Fluid) using 200-μm-deep, 3-mm-wide aluminium planchettes (Wohlwend GmbH). Freeze substitution (FS) was done using the AFS2 machine (Leica microsystems) in a cocktail containing 1% OsO4, 0.1% uranyl acetate and 5% water in acetone. The samples were incubated as follows: 73h at −90°C, temperature increased to −30°C at a rate of 5°C /h. 5h at −30°C, temperature increased to 0°C at a rate of 5°C/h. 4 × 0.5h rinses in water-free acetone at 0°C. This was followed by Spurr resin infiltration (25%—3h at 0°C, 50% overnight at 0°C; 50%—4 h at room temperature; 75%—4 h, 75% overnight, 100% 4 h (×2) and 100% overnight. This was followed by exchange with 100% Spurr resin (4 h × 2, followed by overnight). After this the samples were polymerized in the oven at 60°C for over 2 days. The samples were then cut using an ultramicrotome (Leica UC7) into 70 nm sections and screened by two-dimensional (2D) TEM (Jeol 1400 Flash) to assess sample preparation. For TEM tomography 300 nm sections were poststained with 2% uranyl acetate in 70% methanol (5 min, room temperature) and in Reynolds lead citrate (2 min, room temperature). The tomograms were imaged on a Tecnai F30 microscope using SerialEM and reconstructed using iMOD eTOMO.^38,39^

### FIB-SEM and segmentation

The Spurr resin-embedded samples were prepared as described above. 70 nm sections were collected and screened by 2D TEM (Jeol 1400 Flash) to target cellularizing cells for FIB-SEM imaging as previously described (Ronchi et al., 2021; Mocaer et al., 2023; Shah et al., 2024).^29^ The samples were then mounted on a SEM stub using silver conductive epoxy resin (Ted Pella), gold sputter coated (Quorum Q150RS) and imaged by FIB-SEM. The acquisition was performed using a Crossbeam 540 (Zeiss) following the Atlas 3D Nanotomography workflow. SEM imaging was done with an acceleration voltage of 1.5 kV and a current of 700 pA using an ESB detector (1,100 V grid). Images were acquired at 8 x 8nm2 pixel size and 8 nm slices were removed at each imaging cycle. FIB milling was performed at 1.5 nA current. For image analysis, we binned the dataset to a final voxel size of 16×16×16 nm3. Segmentation and visualization were done with Amira (v.2019.3 or 2020.1; ThermoFisher Scientific).

### Image analysis, quantification and statistical analysis

Image analysis (2D) was done using Fiji.^40,41^ Coenocyte diameter/radius, MT length, nuclear distance from the cortex and PM furrow length from U-ExM images were measured by performing maximum intensity projections of 15-20 sections from the center of the coenocyte. Using the line tool from Fiji, 4 different compartments or nuclei per coenocyte were quantified. MT length (tubulin channel) was measured from the MTOC towards the center of the coenocyte. Nuclear distance from the cortex was calculated as the euclidean distance between the nucleus and the cortical membrane using the DNA channel (Hoechst) and membrane channel (BODIPY) to find the edges of cells in the coenocyte.

### Analysis of timelapse movies with FM4-64 dye

To measure cell diameter and PM furrow length in Figure 1, 3 and 4, we cropped time-lapse movies of FM4-64-labeled coenocytes to have a single coenocyte per movie. For each cropped timelapse, the middle section was used for all the measurements. The line tool from Fiji was used to measure the diameter and PM furrow length manually, for 3-4 PM furrows per coenocyte per treatment. The median value was used for each time point and recorded in Excel. To calculate PM furrow spacing, we selected the time point of 10 min in which PM furrows start to ingress at a value of 2-5 µm. Then a circle line equal to the coenocyte diameter was positioned at the cortex of each coenocyte and defined with a width of 10. Using the plugin “area to line” to create a plot profile with the signal intensity for each furrow, furrow spacing was calculated as the distance between each peak of the plot profile.

Cell perimeter was calculated in a semi-automated manner by first subtracting the background (50 pixels) of each cropped timelapse movie and applying a Gaussian blur filter (sigma=1). Using an Otsu threshold, a black-and-white mask was created and cell perimeter was then measured using the wand-tool and the “measure” function in fiji.

R studio (R software) was used to plot PM furrow length over time, and to change the aesthetics of the plots. Plots were generated in Rstudio. All figures were assembled with Adobe Illustrator.

